# Post-encoding modulation of spatial memory by a GABA_A_-agonist

**DOI:** 10.1101/2021.09.12.458696

**Authors:** Deetje Iggena, Patrizia M. Maier, Sophia M. Häußler, Mario Menk, Heidi Olze, Matthew E. Larkum, Carsten Finke, Christoph J. Ploner

## Abstract

We investigated the role of the post-encoding period for consolidation of self-centered (egocentric) and world-centered (allocentric) spatial memory in neurologically normal human subjects. We used the GABA_A_-ergic anesthetic propofol to transiently modulate neural activity during the early stage of spatial memory consolidation. A total of 52 patients undergoing minor surgery learned to navigate to a target in a five-armed maze derived from animal experiments immediately prior to injection of propofol (early group) or more than 60 minutes before injection (late group). Two hundred and forty minutes after anesthesia, subjects were tested for memory-guided navigation. Our results show a selective impairment of memory-guided navigation in the early group and near-normal performance in the late group. Both egocentric and allocentric navigation were affected, albeit with distinct error patterns. In the egocentric condition, early group patients navigated significantly more often to a wrong alley of the maze but showed normal navigation times, thus suggesting a deficit mainly for memory of sequences of path segments. By contrast, in the allocentric condition, early group patients mostly navigated to the correct alley of the maze but showed a significant increase in detours and prolonged navigation times, thus suggesting a weakened representation of the relationship between landmarks. We conclude that presumably hippocampus-dependent networks contribute to early consolidation of representations underlying both egocentric and allocentric memory-guided navigation. Distinct aspects of these representations are susceptible to GABA_A_-ergic modulation within a post-encoding time-window of less than 60 minutes, indicating a redistribution and reconfiguration of spatial memory networks early during consolidation.

**Significance statement:**

1. Propofol modulates consolidation of spatial representations underlying human spatial navigation.
2. Following administration of propofol, memory-guided navigation using self-centered (egocentric) and world-centered (allocentric) spatial information is impaired.
3. Error patterns after administration of propofol suggest modulation of a post-encoding integration process relevant for ego- and allocentric memory representations.
4. The transient susceptibility of this process to GABA_A_-ergic modulation is consistent with rapid reconfiguration of networks for spatial memory shortly after learning.
5. Propofol provides a pharmacological tool to investigate spatial memory consolidation in humans.

## Introduction

Memory consolidation is an umbrella term for processes that transform novel mental representations into lasting memories over time, including spatial memories (Müller and Pilzecker, 1900; Dudai et al., 2015; Squire et al., 2015). The standard model holds that early memory consolidation is a hippocampus-dependent process on the level of synapses and neural circuits. In contrast, late memory consolidation is conceived as a large-scale rearrangement process characterized by increasing neocortical involvement (Alvarez and Squire, 1994; McClelland et al., 1995). After encoding, synaptic consolidation is thought to occur within seconds to hours while systems consolidation may continue for months or even years (Kelleher et al., 2004; Smith and Squire, 2009; Takashima et al., 2009). Recently, the traditional view on memory consolidation has been challenged by research suggesting that significant hippocampal-neocortical rearrangement can occur within minutes after encoding (Lesburguères et al., 2011; Kitamura et al., 2017; Moon et al., 2020; Tambini and D’Esposito, 2020).

While human lesion studies complement animal research on late memory consolidation, research on early human memory consolidation remains challenging. Few experimental tools exist to transiently interfere with early memory consolidation in both rodents and humans. A potential pharmacological approach is the short-acting anaesthetic propofol (2,6-diisopropylphenol), which is commonly used in routine medical procedures (Sahinovic et al., 2018; Walsh, 2018). Propofol acts as an agonist on the g-aminobutyric-acid (GABA)-A-receptor and as a partial antagonist on (NMDA)-receptors, affecting long-term-potentiation and synaptic consolidation in hippocampal slices as well as hippocampus-dependent memory consolidation (Wei et al., 2002; Nagashima et al., 2005; Zhang et al., 2013; Moon et al., 2020). An fMRI study showed that even sub-hypnotic doses of propofol are sufficient to modulate hippocampal activity in humans (Pryor et al., 2015). Propofol, therefore, represents a promising tool to transiently interfere with hippocampus-dependent cognitive processes and to elucidate how the human brain processes, transforms, and stabilizes memories over time (Vallejo et al., 2019; Moon et al., 2020).

Previous studies showed that formation of spatial memory results from a complex interplay of spatial representations relative to one’s own body coordinates and movements (egocentric representations) with information about spatial relationships in the environment (allocentric representations). During free navigation, both types of information are likely to be acquired simultaneously. It is however unclear whether these representational modes can be clearly dissociated at the neural level (Ekstrom et al., 2014, 2017; Alexander et al., 2020; Ladyka-Wojcik and Barense, 2021). While hippocampus and adjacent regions of the medial temporal lobe have traditionally been associated with allocentric spatial representations, recent research in animals and humans suggests an important role for egocentric memory as well (Kunz et al., 2020; Johnsen and Rytter, 2021; Samanta et al., 2021).

Here, we used propofol to investigate the contribution of hippocampus-dependent networks to early spatial memory consolidation in humans. We reasoned that if the relative contribution of hippocampus-dependent networks to ego- and allocentric spatial representations during the post-encoding period is distinct, this should be reflected in a differential susceptibility to GABA_A_-ergic modulation with propofol. Neurologically normal patients undergoing minor surgery learned to navigate to a target in a virtual five-armed maze. The maze was a modified version of a setup for rodent research that allows to separately investigate ego- and allocentric memory-guided navigation (Rondi-Reig et al., 2006; Iglói et al., 2009). At different time-points after learning to navigate the maze, two groups of patients received a general anesthesia with propofol. After anesthesia, subjects were tested for egocentric and allocentric memory-guided navigation.

## Materials and Methods

### Participants

We included 78 subjects (52 patients, 26 controls; 37 female and 41 male) in our study (Table 1). Patients underwent total intravenous anaesthesia with propofol for minor surgery of nasal septal deviation, sinusitis, or tonsillitis (Table 1). All patients were recruited from the ear-nose-throat-(ENT)-department of the Charité-Universitätsmedizin Berlin during a visit to the outpatient clinic at least one day before surgery. Participants were between 18 and 49 years old, spoke German fluently, had normal or corrected-to-normal vision, normal hearing, reported to be in good health and denied any history of a neuropsychiatric disorder or substance abuse. All participants completed a German version of the Santa Barbara Sense of Direction scale (SBSODs), i.e. a questionnaire that assesses spatial abilities, preferences and experiences (Hegarty et al., 2002).

**Table 1.** Demographic and clinical data of the investigated groups. Data presented as median and interquartile range (25 – 75%).^1^ χ^2^-test, ^2^ K.-Wallis-χ^2^-test

|  | **Control (n=26)** | **Propofol-early (n=26)** | **Propofol-late (n=26)** | ***p-value*** |
| --- | --- | --- | --- | --- |
| Female/male | 15/11 | 13/13 | 9/17 | *0.237*<sup>1</sup> |
| Age | 27.5 (22 – 36.5) | 28 (24 – 32) | 29 (22 – 32) | *0.666*<sup>2</sup> |
| Years of education | 16 (15 – 18) | 15.5 (13 – 18) | 15 (13.75 – 15.5) | *0.245*<sup>2</sup> |
| SBSODs | 5 (4.1 – 5.45) | 4.3 (3.6 – 5.2) | 4.7 (4.3 – 5.1) | *0.294*<sup>2</sup> |
| Medical Procedure | n.a. | Tonsillectomy (n = 8)<br>Rhinoplasty/<br>functional endoscopic<br>sinus surgery (n = 18) | Tonsillectomy (n = 5)<br>Rhinoplasty/<br>functional endoscopic<br>sinus surgery (n = 21) |  |
| Propofol bolus dose (mg) | n.a. | 200 (165 – 200) | 200 (165 – 200) | *0.938* |
| Propofol maintenance dose (mg/kg/h) | n.a. | 6.1 (6 – 6.7) | 6 (6 – 6.5) | *0.841* |
| Duration propofol administration (min) | n.a. | 63.5 (54.3 – 81.3) | 73.5 (55.3 – 98.3) | *0.264* |
| Remifentanil maintenance dose (μg/kg/h) | n.a. | 0.2 (0.2 – 0.21) | 0.2 (0.2 – 0.25) | *0.327* |
| Delay end of learning and propofol (min) | n.a. | 18 (13 – 22.8) | 145 (99.5 – 172) | *< 0.001* |
| Delay end of propofol and recall (min) | n.a. | 246.5 (217.8 – 264.3) | 232 (214.3 – 253.5) | *0.314* |
| Delay end of learning and recall (min) | 301 (282.3 – 341) *a* | 320.5 (297.3 – 356.8) *b* | 441 (413.3 – 510.8) *c* | *< 0.001*<br>*b\*a = 0.180*<br>*b\*c < 0.001*<br>*a\*c < 0.001* |

Twenty-six patients learned a spatial memory task about eighteen minutes before anesthesia (Mdn 18, IQR 13 – 22.8; “propofol-early”; Fig. 1B, Table 1) and 26 patients learned the task about 145 minutes before anesthesia (Mdn 145, IQR 99.5 – 172; “propofol-late”; Fig. 1B, Table 1). Each patient received intravenous general anesthesia, starting with a propofol bolus for anesthesia induction (Mdn 200 mg, IQR 165 - 200), followed by continuous propofol administration with 6mg/kg/h for about 65 minutes for maintenance (Table 1). For analgesia, subjects received a continuous infusion of remifentanil with 0.2μg/kg/min (Table 1). We further recruited 26 age-, sex-, and education-matched healthy control participants via the intranet of the Charité-Universitätsmedizin Berlin (Fig. 1, Table 1). The three subject groups did not differ significantly in terms of age, gender ratio, years of education and SBSODs scores (Table 1).

**Figure 1.**
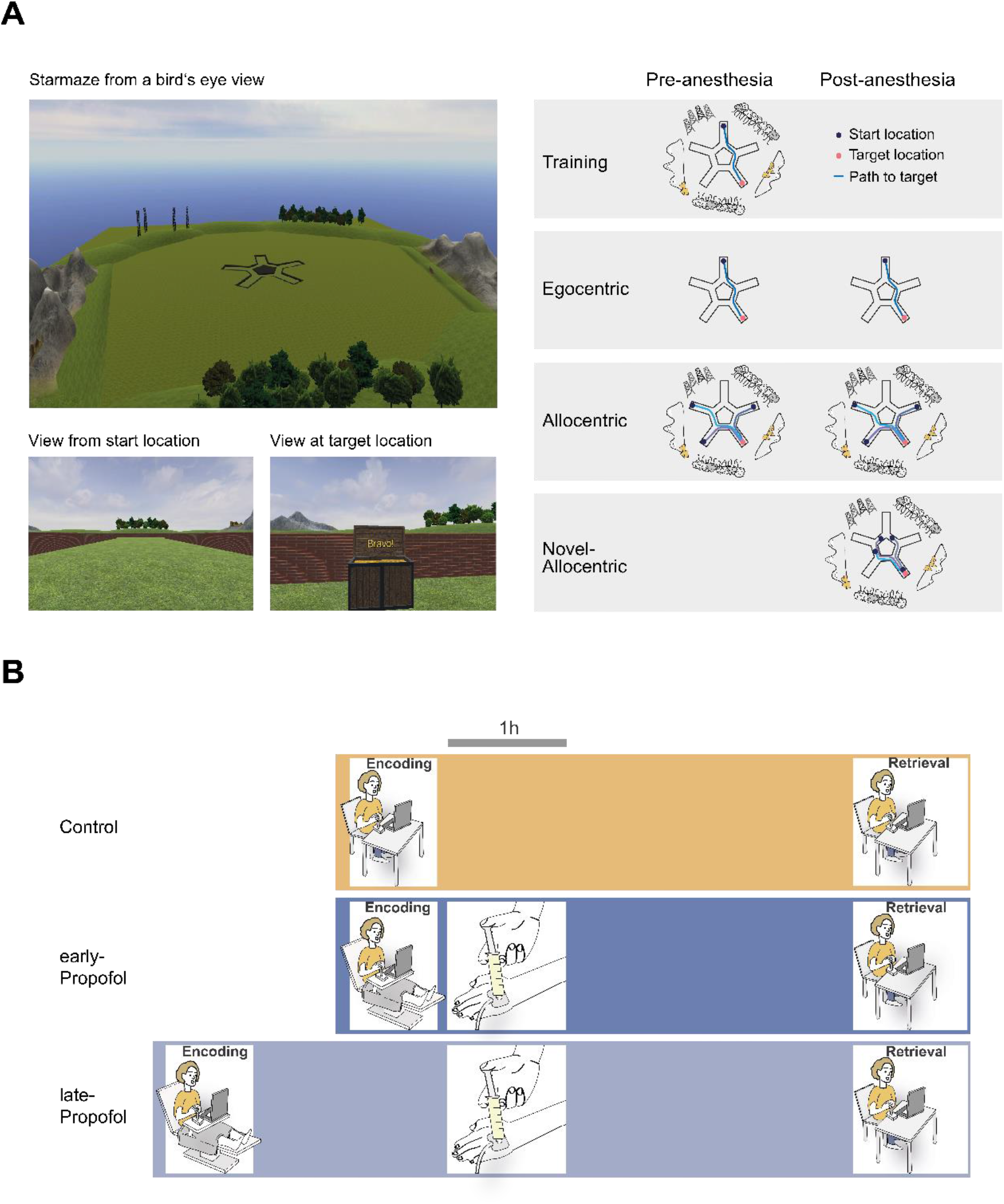
Experimental design. ***A***, Virtual navigation setup. Left, five-armed maze surrounded by environmental cues, bird’s eye view; example views at starting-location and target location in training trials, subject’s view. Right, schematic of the four trial types of the study. Blue lines denote ideal paths connecting starting and target locations. ***B***, Experimental timeline. First row, control group. Second row, early propofol group. Third row, late propofol group. All groups learned to navigate to a target in a virtual maze. The control group received no propofol and was tested about 5h after learning, the early propofol group received propofol 18 minutes after learning and was tested about 5h after learning, the late propofol group received propofol 2.5 h after learning and was tested about 7.5 h after learning.

All participants gave written informed consent. All experimental procedures were conducted according to the declaration of Helsinki and were approved by the local ethics committee of the Charité-Universitätsmedizin Berlin.

## EXPERIMENTAL DESIGN AND STATISTICAL ANALYSES

### Virtual navigation setup

The setup consisted of a virtual star-shaped maze with environmental cues (Fig. 1). The maze was an adapted version of a maze used in previous human and animal studies on navigation (Rondi-Reig et al., 2006; Iglói et al., 2009, 2010). The maze consisted of five symmetrically arranged peripheral alleys connected by five central alleys and was surrounded by five distant environmental cues, embedded in a virtual landscape (2 x forest, 2 x mountains with village, 1 x group of transmission towers; Fig. 1). A treasure was hidden at the end of one of the peripheral alleys (Fig. 1A). The task was presented on a Lenovo Thinkpad X1 Carbon laptop computer (14.0-inch screen). Participants used a joystick controller to move and turn within the environment. We created the stimuli in Blender (version 2.79b, Blender Foundation) and Unity3D (version 2018.2.14f, Unity Technologies). The trial structure and recording of movement trajectories was implemented using the Unity Experiment Framework (Brookes et al., 2020).

### Behavioral testing

Before the experiment, we informed the participants that they would perform a navigation task before and after anesthesia but did not mention any details of the trials in the post-anesthesia testing session. We instructed the participants to search for a treasure hidden somewhere in the virtual maze. The treasure was always at the same position and appeared as soon as the subject reached its location (training trial). We also informed participants that in some trials they would have to indicate the memorized position of the treasure by pressing a red button as soon as they had reached its location (probe trial). In these trials, the treasure would not appear, even if the location was correctly remembered. We asked the participants to navigate on a direct path to the treasure and informed them that neither the maze nor the environment would change during the experiment. Trials were terminated four seconds after the treasure appeared (training trials) or after the button was pressed (probe trials). If neither event occurred within ninety seconds, the trial was terminated. Prior to the starmaze task, all participants familiarized themselves with the joystick and task requirements by completing three practise trials in a simple virtual three-armed maze.

### Pre-anesthesia session

The pre-anesthesia session lasted about 15 minutes and aimed to ensure encoding of both egocentric and allocentric spatial representations (Fig. 1A). During the first four training-trials, participants navigated freely until they found the treasure. For navigation to the remembered location of the treasure, they could either reproduce their own successful path from a previous trial (egocentric strategy) and/or orient themselves based on the spatial relationships between environmental cues and the location of the treasure (allocentric strategy) (Iglói et al., 2009, 2010). After four training trials, participants had to indicate where the treasure was hidden in one probe trial. In case the participants failed to locate the target correctly, they were allowed one more training and one more probe trial. We then removed all environmental cues for the egocentric condition and participants had to navigate from the original starting point to the treasure. This manipulation was chosen to enforce egocentric sequence-based navigation (Iglói et al., 2009, 2010). After three training trials, participants had to indicate where the treasure was hidden in one probe trial. Afterwards, the environmental cues reappeared for the allocentric condition, and we informed the participants that their starting point would vary between trials. This manipulation was chosen to enforce allocentric landmark-based navigation (Iglói et al., 2009, 2010). After six training trials, participants had to indicate where the treasure was hidden in three consecutive probe trials. We assumed proper learning of the task if participants solved the probe trial requiring egocentric navigation and at least two out of three probe trials requiring allocentric navigation.

### Post-anesthesia session

The post-anesthesia session lasted about 15 minutes and aimed to test memory of spatial representations from the pre-anesthesia session. Before the session, we informed the participants that they would not receive feedback throughout the session and would always have to indicate the location of the treasure (i.e., probe trials only). Testing consisted of three experimental trial types. To test memory of spatial representations for egocentric navigation, we removed all environmental cues for the first three trials (egocentric trials). In the following trials, all environmental cues were always visible. To test for retention of spatial representations for allocentric navigation, participants started from varying starting locations for seven consecutive trials. The first three locations had also been used during the pre-anesthesia session (allocentric trials), the last four were novel starting locations (novel allocentric trials). These latter trials were intended to test for flexible landmark-based navigation.

### Data acquisition and statistical analysis

During virtual navigation, we recorded virtual positions within the maze as x- and y-coordinates in a Cartesian coordinate system combined with a timestamp at an average sampling rate of 110 Hz. For analysis, we determined whether participants successfully solved the task by calculating the final distance to the target location (Eq.: Σ(√[(x(treasure) – x(end))^2 – (y(treasure) – y(end))^2])). A trial was considered successful when the final distance to the target location was less than 0.1 virtual meter.

To assess and identify changes in successful navigational behavior, we first extracted the trial-duration from the timestamps (Eq.: t(end) - t(1)) to calculate the navigation time in successful trials. Second, we calculated the path length (Eq.: Σ(√[(x(i + 1) – x(i))^2 – (y(i + 1) – y(i))^2])) to determine the total distance covered to reach the target location in successful trials. Third, we calculated the distance to the target location averaged across all time stamps of a trial (Eq.: Σ(√[(x(treasure) – x(i))^2 – (y(treasure) – y(i))^2])/length(coordinates)) to reveal changes in navigational behavior that are not immediately evident in a mere analysis of path length. The average distance to target is a measure of target proximity and functions a surrogate for the degree of uncertainty in navigational behavior. To account for alternating starting positions, the path-length and the average distance to target were normalized by calculating the absolute percent error (path-error/ distance-error = (ideal value – actual value)/ ideal value) * 100. To avoid artificial error increases when participants started within 0.1 virtual meters of the target and the absolute path length did not exceed 0.1 virtual meters, we set both errors to zero. Since we were primarily interested in performance changes between pre-anesthesia and post-anesthesia sessions, we subtracted the first measured value from the second measured value to receive a delta score (Eq.: Δ = value 2 – value 1) for all main variables (Dimitrov and Rumrill, 2003). All analyses of navigational behavior were performed in Matlab (Matlab 2020b, Mathworks, USA).

All statistical analyses were performed in R (v. 4.1.0). Shapiro-Wilk-testing showed that the assumption of normality had to be rejected for our main dependent variables. We thus chose non-parametric statistical tests for statistical analyses. To detect between-group-differences of metric variables, we used the non-parametric Kruskal-Wallis-test. Mann-Whitney-Tests with Bonferroni correction were used for subsequent post-hoc comparisons between groups. For all statistical tests, the significance level was set to 0.05.

### Code accessibility

Analysis-script and functions are available at github: https://github.com/DeetieIggena/five-arm-maze-analysis/

## Results

### Egocentric navigation

In pre-anesthesia egocentric trials, all participants successfully learned the sequence of path segments and left and right turns to reach the target location (Fig. 2). Navigation behavior in the pre-anesthesia session did not differ between groups (time: K.-Wallis-χ^2^(2) = 0.802, p < 0.670; path error: K.-Wallis-χ^2^(2) = 1.05, p = 0.592; distance error: K.-Wallis-χ^2^(2) = 1.458, p = 0.482).

**Figure 2.**
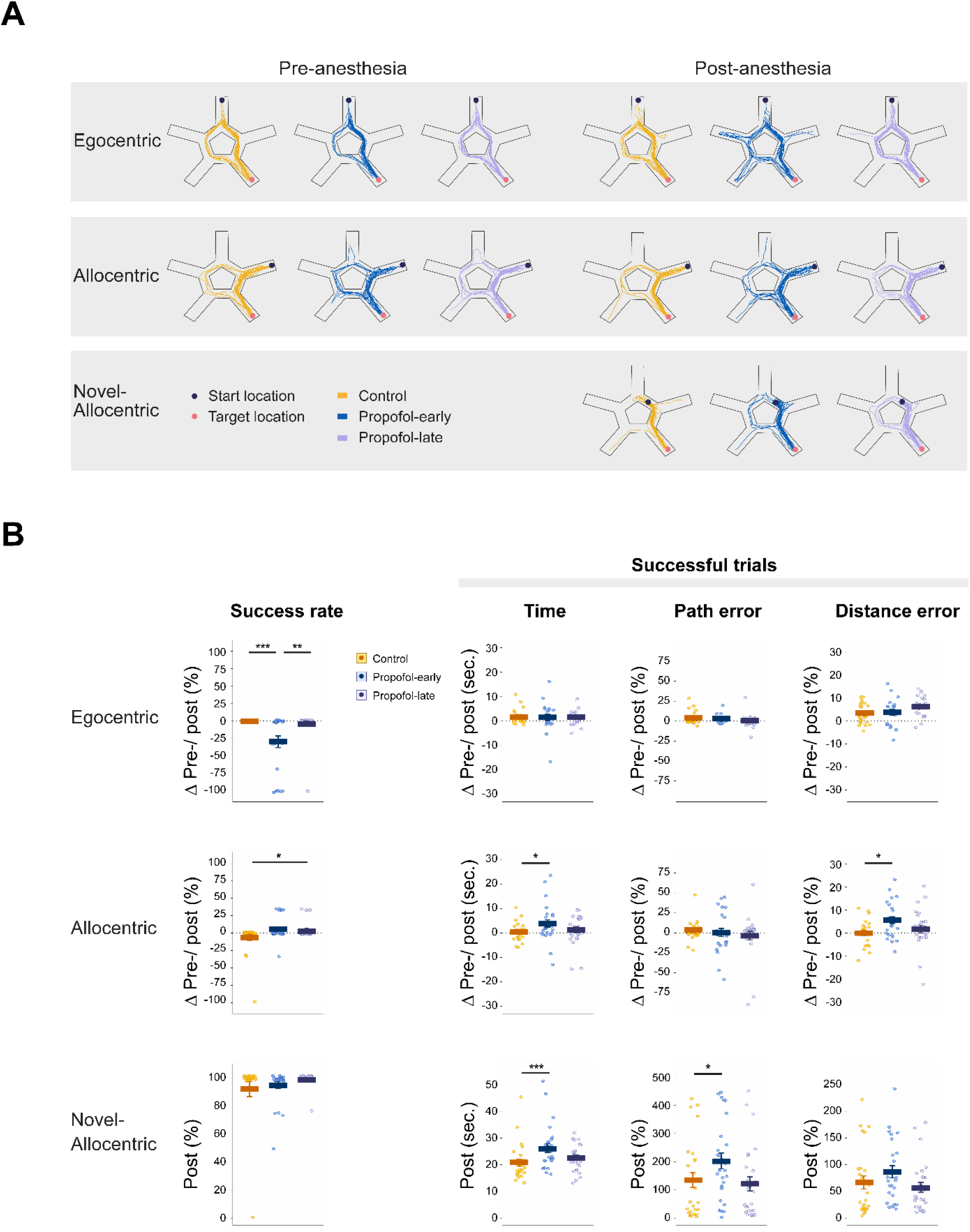
Navigation performance. ***A***, Exemplary navigation paths from the three investigated subject groups. Left, pre-anesthesia paths; right, post-anesthesia paths. First row, egocentric condition, second row allocentric condition, third row novel-allocentric condition. Note increased number of paths into wrong alleys in the propofol-early group in the egocentric condition. Note increased number of detours in the propofol-early group in the novel-allocentric condition. ***B***, Group performance. First row, egocentric condition, second row allocentric condition, third row novel-allocentric condition. Note that in the egocentric condition, administration of propofol caused a significant and selective pre-/postanesthesia decrease in the number of successful trials. Note that in the allocentric condition, propofol caused a pre-/postanesthesia increase in navigation time and distance-error in the propofol-early group. Note that in the novel-allocentric condition (post-anesthesia testing only), administration of propofol caused an increase in navigation times and path error in the propofol-early group compared to the other groups. Yellow, control group; blue, propofol-early group, violet propofol-late group. Data presented as means and single datapoints for each participant. \**p* < 0.05; \*\**p* < 0.01; \*\*\**p* < 0.001.

At post-anesthesia testing, the propofol-early group showed a 29.5% decrease in successful egocentric trials, i.e. subjects terminated navigation in a wrong alley of the maze more frequently, whereas the success rate of the control group did not change and the success rate of the propofol-late group decreased by 4% only (K.-Wallis-χ^2^(2) = 18.624, p < 0.001; control vs. propofol-early, p = 0.002, control vs. propofol-late, p = 1.0, propofol-early vs. propofol-late, p = 0.01. Fig. 2). However, other parameters of egocentric navigation remained unchanged in both propofol groups. Propofol caused no significant differences of pre-/post-anesthesia changes in navigation time (K.-Wallis-χ^2^(2) = 0.115, p = 0.944), path error (K.-Wallis-χ^2^(2) = 0.603, p = 0.740), and distance error (K.- Wallis-χ^2^(2) = 5.908, p = 0.052) between groups. Thus, while the control and the late-propofol groups correctly reproduced the paths learned in the pre-anesthesia session, the early-propofol group more frequently generated erroneous paths without changes in navigational efficiency or signs of uncertainty, therefore suggesting a deficit mainly in memory of sequences of path segments.

### Allocentric navigation – repeated starting positions

In pre-anesthesia allocentric trials, all participants successfully learned to reach the target location by using environmental landmarks (Fig. 2). Navigation behavior in the pre-anesthesia session did not differ between groups (time: K.-Wallis-χ^2^(2) = 2.569, p = 0.277; path-error: K.-Wallis-χ^2^(2) = 0.472, p = 0.790; distance-error: K.- Wallis-χ^2^(2) = 3.330, p = 0.189).

At post-anesthesia testing, when participants started from positions that also been used in the pre-anesthesia session, we found minor changes in success rates between groups that fell far below the changes seen in egocentric trials with pre-/post-anesthesia changes of up to 6.5 % in the propofol-late group (K.-Wallis-χ^2^(2) = 7.190, p = 0.027; control vs. propofol-early, p = 0.063, control vs. propofol-late, p = 0.048, propofol-early vs. propofol-late, p = 1.0. Fig. 2). However, propofol induced significant changes in further navigational parameters in the propofol-early group. We found a significant pre-/post-anesthesia increase in navigation time of 3.9 s compared to 0.3 s in controls (K.-Wallis-χ^2^(2) = 6.429, p = 0.040; control vs. propofol-early, p = 0.033, control vs. propofol-late, p = 0.436, propofol-early vs. propofol-late, p = 0.912, Fig. 2B). Apparently, subjects in the propofol-early group spent more time at maze locations remote from the target, as also reflected by a 5.6% pre-/post-anesthesia increase in distance error (K.-Wallis-χ^2^(2) = 7.594, p = 0.022; control vs. propofol-early, p = 0.017, control vs. propofol-late, p = 1.0, propofol-early vs. propofol-late, p = 0.241. Fig. 2). However, the change in the temporal properties of navigational behavior in the propofol-early group was not accompanied by significant modifications of path geometry, as indicated by similar pre-/post-anesthesia changes in path-error between groups (K.-Wallis-χ^2^(2) = 1.045, p = 0.593). These results therefore suggest that propofol did not significantly alter recognition of landmarks or memory of the paths that had been travelled in the pre-anesthesia session but rather induced a significant increase in navigational uncertainty.

### Allocentric navigation – novel starting positions

In the post-anesthesia session, subjects started from four additional locations in the allocentric condition that were not used in the pre-anesthesia session. Successful navigation thus critically depended on the ability to flexibly update subject position in relation to environmental landmarks rather than on mere repetition of navigational paths that had been travelled in the pre-anesthesia session (Fig. 1). Consistent with our previous observation in allocentric trials with repeated starting positions, we only observed few unsuccessful trials in each group with a success rate of almost 100% in each group (K.-Wallis-χ^2^(2) = 5.516, p = 0.063; Fig. 2).

However, subjects in the propofol-early group showed significantly longer navigation time to reach the target than the other two groups (K.-Wallis-χ^2^(2) = 16.698, p < 0.001; propofol-early vs. control, p < 0.001, propofol-late vs. control, p = 0.051, propofol-early vs. propofol-late, p = 0.187; Figure 2). We therefore analysed the number of entries into different maze zones and calculated the time spent within in these zones. This analysis showed that participants in the propofol-early group visited incorrect external alleys of the maze significantly more often than controls (K.-Wallis-χ^2^(2) = 8.008, p = 0.018; propofol-early vs. control, p 0.017, propofol-late vs. control, p = 0.058, propofol-early vs. propofol-late, p = 1.0) and subsequently spent significantly more time within these incorrect alleys than the control group. The propofol-late group only showed a trend for these parameters (K.-Wallis-χ^2^(2) = 8.008, p = 0.018; propofol-early vs. control, p = 0.013, propofol-late vs. control, p = 0.060, propofol-early vs. propofol-late, p = 1.0). The increase in navigation time in the propofol-early group can further be explained by an increase in path-length as reflected in an average path-error of 197.5%, compared to 121.3% in the control group and 122.2% in the propofol-late group (K.-Wallis-χ^2^(2) = 9.525, p = 0.009; early-propofol vs. control, p = 0.012, late-propofol vs. control, p = 1.0, early-propofol vs. late-propofol, p = 0.056, Figure 2). However, there was no evidence that participants of the early-propofol group spent more time at a greater distance from the target, as the distance-error did not vary between groups (K.-Wallis-χ^2^(2) = 4.444, p = 0.108, Figure 2). Thus, when participants were forced to base navigation solely on a flexible integration of their navigational position with the spatial relationship of landmarks in the post-anesthesia session, the propofol-early group took detours and spent more time in incorrect alleys with concomittant increases in navigation time and path-error. These results therefore suggest a weakened representation of the spatial relationship between landmarks in the propofol-early group.

## Discussion

We investigated effects of the GABA_A_-ergic anesthetic propofol on consolidation of spatial memory in humans. We used a virtual variant of a star-shaped maze that allows to disentangle ego- and allocentric navigational strategies and that has previously been shown to be sensitive to hippocampal dysfunction (Rondi-Reig et al., 2006; Iglói et al., 2009). Our results show that administration of propofol after learning to navigate the maze significantly impairs representations required for later memory-guided navigation. These effects were confined to a brief time window of less than one hour after learning and affected retrieval conditions that prompted the use of egocentric as well as allocentric strategies. Different error patterns between conditions however suggest that propofol affected consolidation of distinct aspects of egocentric and allocentric spatial representations. Memory-guided navigation performance in the egocentric condition was consistent with impaired representation of sequences of path segments whereas navigation performance in allocentric conditions mainly suggested a weakened representation of the spatial relationship of landmarks. The transientness of these effects further implies early reconfiguration of hippocampus-dependent networks during spatial memory consolidation.

In our subjects, propofol was administered systemically. With a context-sensitive half-time of less than 10 minutes (Hughes et al., 1992; Sahinovic et al., 2018) its effects are quite selective in time but necessarily not confined to a distinct region of the brain. However, it has been shown previously that propofol modulates neuronal networks by activating intra- and extra-synaptic GABA_A_ receptors. The influx of anions attenuates synaptic transmission (Collins, 1988; Otsuka et al., 1992; Orser et al., 1994), regulates LTP (Wang et al., 2006) and disturbs the rhythmic activity of neurons (Perouansky and Pearce, 2011). GABA_A_ receptor subtypes are unequally sensitive to propofol (Wang et al., 2018) and unevenly distributed throughout the CNS (Fritschy et al., 1997; Pirker et al., 2000). GABA_A_ receptors containing the α5-subunit (α5GABA_A_) are probably crucial for memory effects of propofol (Perouansky and Pearce, 2011; Engin et al., 2020). α5GABA_A_ receptors are highly expressed in the hippocampus, where this specific subtype contributes to 25% of all GABA_A_ receptors (Fritschy et al., 1997; Pirker et al., 2000). Outside the hippocampal formation, the relative prevalence of α5GABA_A_-receptors is less than 5% (Fritschy et al., 1997). Pharmacological stimulation of hippocampal α5GABA_A_ receptors has been shown to reduce hippocampal excitability, to disturb hippocampal sharp wave and ripple oscillations and to prevent excessive activation of excitatory synapses by downregulating LTP (Papatheodoropoulos and Koniaris, 2011; Viereckel et al., 2013; Davenport et al., 2021).

Consistent with the anatomical distribution of α5GABA_A_-receptors, electrophysiological experiments showed that propofol modulates key mechanisms of memory consolidation. In hippocampal slices, propofol has been shown to transiently affect induction of LTP (Nagashima et al., 2005; Takamatsu et al., 2005) and maintenance of LTP in the CA1 region of the rat hippocampus (Wei et al., 2002). In line with these findings, propofol infusion during memory consolidation impaired hippocampus-dependent spatial memory in rats and decreased recall performance in a word-list task relying on hippocampal integrity in humans (Zhang et al., 2013; Moon et al., 2020). Furthermore, a PET-study detected reduced hippocampal glucose-metabolism during propofol administration (Sun et al., 2008) and fMRI experiments showed suppressed hippocampal activity during encoding of emotional pictures under continuous propofol infusion in sub-anesthetic doses (Pryor et al., 2015). It appears thus likely that the memory effects of propofol observed here are largely mediated by hippocampus-dependent neural networks.

Most theories address the mechanisms underlying memory consolidation either on a local synaptic level or on a large-scale (i.e. hippocampal-neocortical) systems level. These two groups of mechanisms are frequently considered to be associated with distinct time scales (Alvarez and Squire, 1994; McClelland et al., 1995). Synaptic consolidation is thought to operate for up to some hours, whereas systems consolidation may continue for years (Dudai et al., 2015; Squire et al., 2015). However, recent evidence shows that processes of systems level consolidation can already be observed in the immediate post-encoding period (Dudai et al., 2015; Tambini and Davachi, 2019). Reactivation of neural activity in the hippocampus in the post-encoding period has been shown to be associated with activity across hippocampal-neocortical networks that determines later recall. For example, functional connectivity between the hippocampus and the lateral occipital complex during some minutes following encoding in a visual associative memory task correlated with later memory performance (Tambini et al., 2010). Likewise, multi-voxel activity patterns in the human medial temporal lobe and retrosplenial cortex during encoding and the immediate post-encoding period have been shown to predict later recall in a similar task (Staresina et al., 2013). In a recent study, interference with hippocampal-neocortical interactions by transcranial magnetic stimulation over neocortex in a time window of up to 50 minutes following encoding of object-face associations led to a selective deficit in associative memory while item memory for objects and faces was spared (Tambini and D’Esposito, 2020). Our results complement these findings by providing further evidence that modulation of post-encoding neural activity in a time window of less than one hour can causally affect systems consolidation, i.e. on a timescale that matches processes of synaptic memory consolidation (Dudai et al., 2015). Moreover, the pattern of impaired sequences of path segments and spatial relationships of landmarks with preserved path segments and landmarks is consistent with the hypothesis that neural activity in the early consolidation period not only strengthens individual memories, but also reflects an ongoing integration process that binds distinct items and distributes memory representations across hippocampal-neocortical networks (Tambini and Davachi, 2019).

Like in previous studies in animal models and humans, we used different conditions at retrieval to prompt the use of distinct strategies for memory-guided navigation. Removal of all landmarks was intended to force subjects to rely on representations of sequences of path segments that had been travelled in the pre-anesthesia session (egocentric trials). Shifting of starting points with landmarks present was intended to allow for additional orienting by using the spatial relationships between landmarks (allocentric trials) or to force subjects to solely rely on these relationships (novel allocentric trials). However, during natural behavior, these representational modes are almost always used in combination, with their relative contribution depending on individual preferences, abilities and contextual factors (Ekstrom et al., 2014, 2017; Johnsen and Rytter, 2021; Ladyka-Wojcik and Barense, 2021). While several distinct local neuronal populations in hippocampus and entorhinal cortex code spatial, temporal and visual features required for memory-guided navigation (Moser et al., 2015; Eichenbaum, 2017), recent evidence suggests that overlapping groups of neurons support allocentric as well as egocentric spatial representations (Alexander et al., 2020). Similarly, on the level of large-scale networks, largely overlapping brain regions including hippocampus, entorhinal cortex, parietal cortex, retrosplenial cortex and others may be involved in shared processing of information for egocentric and allocentric representations (Chrastil, 2013; Ekstrom et al., 2017). By this view, both representational modes may represent a continuum supported by an extended network with changing hubs rather than by clearly separable neural substrates.

Supporting the hypothesis of shared neural resources for egocentric and allocentric processing, rodent experiments employing the star maze task demonstrated that both sequence- and landmark-based navigation rely on the integrity of hippocampal subregion CA1 (Rondi-Reig et al., 2006). Subsequent fMRI-experiments in humans with a highly similar task showed that sequence-based navigation activates the left and landmark-based navigation activates the right hippocampus (Iglói et al., 2010). The impairments observed here fit these observations and further suggest that hippocampus-dependent networks provide computations relevant to early consolidation of allocentric as well as egocentric memory representations (Kunz et al., 2020; Johnsen and Rytter, 2021; Samanta et al., 2021). Alternatively, at least in our task, administration of propofol may have affected a central spatio-temporal binding process that yields seemingly distinct behavioral manifestations - depending on the particular navigational context imposed by the experimenter. A common denominator behind the different deficits in our task may thus be a central impairment in associating navigational information in time (egocentric condition) and space (allocentric conditions). Confirmation of this hypothesis will however require a combination of our neuro-pharmacological approach with electrophysiological recordings in experimental animals.

Taken together, our results provide additional evidence for a significant role of the post-encoding period for later recall (Dudai et al., 2015; Tambini and Davachi, 2019). Our findings are consistent with the hypothesis that neural activity in a brief time window following learning is more than the persistence of a comparatively punctual encoding process. In this case, a more global deterioration of navigational parameters at retrieval would have been expected. Rather, the selectivity of the deficits points to a time-limited associative process that is necessary for the formation of integrated spatial representations that are critical for later memory-guided navigation - in particular those that require sequencing of path segments and flexible use of spatial relationships between landmarks. Our approach only allows for limited inferences on involved neural substrates. However, the known distribution of GABA-receptors together with previous behavioral and imaging evidence of propofol effects on the human brain (Pryor et al., 2015; Moon et al., 2020) make rapidly reconfigurating hippocampus-dependent networks a likely candidate for this process.

## Acknowledgements

This study was funded by the Deutsche Forschungsgemeinschaft (DFG, German Research Foundation) – Project number 327654276 – SFB 1315.

We thank the study participants and the staff of the Department of Otorhinolaryngology and the Department of Anesthesiology for their kind support of our study.

## Author contributions

D.I., P.M.M., C.F. and C.J.P. designed research; D.I. and P.M.M. performed research; S.H., M.M. and H.O. contributed unpublished reagents/analytic tools; D.I., P.M.M., M.E.L., C.F. and C.J.P. analyzed data and wrote the paper.

## References

Alexander AS, Robinson JC, Dannenberg H, Kinsky NR, Levy SJ, Mau W, Chapman GW, Sullivan DW, Hasselmo ME (2020) Neurophysiological coding of space and time in the hippocampus, entorhinal cortex, and retrosplenial cortex. Brain Neurosci Adv 4:1–18.

Alvarez P, Squire LR (1994) Memory consolidation and the medial temporal lobe: A simple network model. Proc Natl Acad Sci 91:7041–7045.

Brookes J, Warburton M, Alghadier M, Mon-Williams M, Mushtaq F (2020) Studying human behavior with virtual reality: The Unity Experiment Framework. Behav Res Methods 52:455–463.

Chrastil ER (2013) Neural evidence supports a novel framework for spatial navigation. Psychon Bull Rev 20:208–227.

Collins GGS (1988) Effects of the anaesthetic 2,6-diisopropylphenol on synaptic transmission in the rat olfactory cortex slice. Br J Pharmacol 95:939–949.

Davenport CM, Rajappa R, Katchan L, Taylor CR, Tsai M-C, Smith CM, de Jong JW, Arnold DB, Lammel S, Kramer RH (2021) Relocation of an extrasynaptic GABA_A_ receptor to inhibitory synapses freezes excitatory synaptic strength and preserves memory. Neuron 109:123–134.

Dimitrov DM, Rumrill PD (2003) Pretest-posttest designs and measurement of change. Work 20:159–165.

Dudai Y, Karni A, Born J (2015) The consolidation and transformation of memory. Neuron 88:20–32.

Eichenbaum H (2017) On the integration of space, time, and memory. Neuron 95:1007–1018.

Ekstrom AD, Arnold AEGF, Iaria G (2014) A critical review of the allocentric spatial representation and its neural underpinnings: Toward a network-based perspective. Front Hum Neurosci 8:1–15.

Ekstrom AD, Huffman DJ, Starrett M (2017) Where Are You Going? The Neurobiology of Navigation. J Neurophysiol 118:3328–3344.

Engin E, Sigal M, Benke D, Zeller A, Rudolph U (2020) Bidirectional regulation of distinct memory domains by α5-subunit-containing GABA_A_ receptors in CA1 pyramidal neurons. Learn Mem 27:423–428.

Fritschy JM, Benke D, Johnson DK, Mohler H, Rudolph U (1997) GABA_A_-receptor α-subunit is an essential prerequisite for receptor formation in vivo. Neuroscience 81:1043–1053.

Hegarty M, Richardson AE, Montello DR, Lovelace K, Subbiah I (2002) Development of a self-report measure of environmental spatial ability. Intelligence 30:425–447.

Hughes MA, Glass PS, Jacobs JR (1992) Context-sensitive half-time in multicompartment pharmacokinetic models for intravenous anesthetic drugs. Anesthesiology 76:334–341.

Iglói K, Doeller CF, Berthoz A, Rondi-Reig L, Burgess N (2010) Lateralized human hippocampal activity predicts navigation based on sequence or place memory. Proc Natl Acad Sci 107:14466–14471.

Iglói K, Zaoui M, Berthoz A, Rondi-Reig L (2009) Sequential egocentric strategy is acquired as early as allocentric strategy: Parallel acquisition of these two navigation strategies. Hippocampus 19:1199–1211.

Johnsen SHW, Rytter HM (2021) Dissociating spatial strategies in animal research: Critical methodological review with focus on egocentric navigation and the hippocampus. Neurosci Biobehav Rev 126:57–78.

Kelleher RJ, Govindarajan A, Tonegawa S (2004) Translational regulatory mechanisms in persistent forms of synaptic plasticity. Neuron 44:59–73.

Kitamura T, Ogawa SK, Roy DS, Okuyama T, Morrissey MD, Smith LM, Redondo RL, Tonegawa S (2017) Engrams and circuits crucial for systems consolidation of a memory. Science 356:73–78.

Kunz L, Brandt A, Reinacher P, Staresina B, Reifenstein E, Weidemann C, Herweg N, Tsitsiklis M, Kempter R, Kahana M, Schulze-Bonhage A, Jacobs J (2020) A neural code for egocentric spatial maps in the human medial temporal lobe. Neuron 109:2781–2796.

Ladyka-Wojcik N, Barense MD (2021) Reframing spatial frames of reference: What can aging tell us about egocentric and allocentric navigation? Wiley Interdiscip Rev Cogn Sci 12:1–12.

Lesburguères E, Gobbo OL, Alaux-Cantin S, Hambucken A, Trifilieff P, Bontempi B (2011) Early tagging of cortical networks is required for the formation of enduring associative memory. Science 331:924–928.

McClelland JL, McNaughton BL, O’Reilly RC (1995) Why there are complementary learning systems in the hippocampus and neocortex: Insights from the successes and failures of connectionist models of learning and memory. Psychol Rev 102:419–457.

Moon DU, Esfahani-Bayerl N, Finke C, Salchow DJ, Menk M, Bayerl S, Kempter R, Ploner CJ (2020) Propofol modulates early memory consolidation in humans. eNeuro 7:1–9.

Moser M-B, Rowland DC, Moser EI (2015) Place cells, grid cells, and memory. Cold Spring Harb Perspect Biol 7:1–15.

Müller GE, Pilzecker A (1900) Experimentelle Beiträge zur Lehre vom Gedächtnis. Z Psych u Physiol d Sinnesorgane 1:1–300.

Nagashima K, Zorumski CF, Izumi Y (2005) Propofol inhibits long-term potentiation but not long-term depression in rat hippocampal slices. Anesthesiology 103:318–326.

Orser BA, Wang LY, Pennefather PS, MacDonald JF (1994) Propofol modulates activation and desensitization of GABA_A_ receptors in cultured murine hippocampal neurons. J Neurosci 14:7747–7760.

Otsuka H, Yamamura T, Hanaoka Y, Kemmotsu O (1992) Does propofol enhance GABA-mediated inhibition? J Anesth 6:305–311.

Papatheodoropoulos C, Koniaris E (2011) α5GABA_A_ receptors regulate hippocampal sharp wave-ripple activity in vitro. Neuropharmacology 60:662–673.

Perouansky M, Pearce RA (2011) How we recall (or don’t): The hippocampal memory machine and anesthetic amnesia. Can J Anesth 58:157–166.

Pirker S, Schwarzer C, Wieselthaler A, Sieghart W, Sperk G (2000) GABA_A_ receptors: Immunocytochemical distribution of 13 subunits in the adult rat brain. Neuroscience 101:815–850.

Pryor KO, Root JC, Mehta M, Stern E, Pan H, Veselis RA, Silbersweig DA (2015) Effect of propofol on the medial temporal lobe emotional memory system: A functional magnetic resonance imaging study in human subjects. Br J Anaesth 115:104–113.

Rondi-Reig L, Petit GH, Tobin C, Tonegawa S, Mariani J, Berthoz A (2006) Impaired sequential egocentric and allocentric memories in forebrain-specific-NMDA receptor knock-out mice during a new task dissociating strategies of navigation. J Neurosci 26:4071–4081.

Sahinovic MM, Struys MMRF, Absalom AR (2018) Clinical pharmacokinetics and pharmacodynamics of propofol. Clin Pharmacokinet 57:1539–1558.

Samanta A, van Rongen LS, Rossato JI, Jacobse J, Schoenfeld R, Genzel L (2021) Sleep leads to brain-wide neural changes independent of allocentric and egocentric spatial training in humans and rats. Cereb Cortex 0:1–16.

Smith CN, Squire LR (2009) Medial temporal lobe activity during retrieval of semantic memory is related to the age of the memory. J Neurosci 29:930–938.

Squire LR, Genzel L, Wixted JT, Morris RG (2015) Memory consolidation. Cold Spring Harb Perspect Biol 7:1–21.

Staresina BP, Alink A, Kriegeskorte N, Henson RN (2013) Awake reactivation predicts memory in humans. Proc Natl Acad Sci 110:21159–21164.

Sun X, Zhang H, Gao C, Zhang G, Xu L, Lv M, Chai W (2008) Imaging the effects of propofol on human cerebral glucose metabolism using positron emission tomography. J Int Med Res 36:1305–1310.

Takamatsu I, Sekiguchi M, Wada K, Sato T, Ozaki M (2005) Propofol-mediated impairment of CA1 long-term potentiation in mouse hippocampal slices. Neurosci Lett 389:129–132.

Takashima A, Nieuwenhuis ILC, Jensen O, Talamini LM, Rijpkema M, Fernández G (2009) Shift from hippocampal to neocortical centered retrieval network with consolidation. J Neurosci 29:10087–10093.

Tambini A, D’Esposito M (2020) Causal contribution of awake post-encoding processes to episodic memory consolidation. Curr Biol 30:3533–3543.

Tambini A, Davachi L (2019) Awake reactivation of prior experiences consolidates memories and biases cognition. Trends Cogn Sci 23:876–890.

Tambini A, Ketz N, Davachi L (2010) Enhanced brain correlations during rest are related to memory for recent experiences. Neuron 65:280–290.

Vallejo AG, Kroes MCW, Rey E, Acedo MV, Moratti S, Fernández G, Strange BA (2019) Propofol-induced deep sedation reduces emotional episodic memory reconsolidation in humans. Sci Adv 5:1–8.

Viereckel T, Kostic M, Bähner F, Draguhn A, Both M (2013) Effects of the GABA-uptake blocker NNC-711 on spontaneous sharp wave-ripple complexes in mouse hippocampal slices. Hippocampus 23:323–329.

Walsh CT (2018) Propofol: Milk of amnesia. Cell 175:10–13.

Wang B, Lv K, Liu H, Su Y, Wang H, Wang S, Bao S, Zhou WH, Lian QQ (2018) Contribution of the α5GABA_A_ receptor to the discriminative stimulus effects of propofol in rat. Neuroreport 29:347–352.

Wang W, Wang H, Gong N, Xu T-L (2006) Changes of K+-Cl-cotransporter 2 (KCC2) and circuit activity in propofol-induced impairment of long-term potentiation in rat hippocampal slices. Brain Res Bull 70:444–449.

Wei H, Xiong W, Yang S, Zhou Q, Liang C, Zeng BX, Xu L (2002) Propofol facilitates the development of long-term depression (LTD) and impairs the maintenance of long-term potentiation (LTP) in the CA1 region of the hippocampus of anesthetized rats. Neurosci Lett 324:181–184.

Zhang J, Zhang X, Jiang W (2013) Propofol impairs spatial memory consolidation and prevents learning-induced increase in hippocampal matrix metalloproteinase-9 levels in rat. Neuroreport 24:831–836.

